# Wild boar trade and African swine fever risk of introduction into new territories: A quantitative release assessment with retrospective data of wild boar shipments to France and Spain (2010-2017)

**DOI:** 10.1101/2025.08.02.668282

**Authors:** F. Jori, D.R.J. Pleydell, E. Burnichon, J. Casal, J.A. Barasona

**Affiliations:** UMR ASTRE (Animal, Health, Territories, Risks and Ecosystems), CIRAD, INRAE, Campus International de Baillarguet, Montpellier, 34398, France; Faculty of Veterinary Science, Universitat Autònoma de Barcelona, Barcelona, Spain; VISAVET, Animal Health Surveillance Center, Madrid, Spain; Animal Health Department, Faculty of Veterinary, Complutense University of Madrid, Madrid, Spain

**Keywords:** African swine fever, *Sus scrofa*, trade, export, import, European Union, assessment, sport hunting

## Abstract

Wild boar (WB) (*Sus scrofa*) farming and trade in EU countries consistently developed at the end of the last century, primarily for repopulating hunting estates within the sport hunting industry. Since the introduction of African swine fever (ASF) in the Caucasus in 2007 and its subsequent spread from Eastern to Central Europe, international WB trade increased the risk of ASF introduction into new territories. In order to prevent such events, the EU veterinary authorities decided to ban WB trade in June 2018. In our study, we documented retrospective WB trade movements to France and Spain between 2010 and 2017, and used this data to undertake a quantitative risk assessment for ASF introduction resulting from those shipments. A total of 127 trade events introducing live WB into France (n=93) and Spain (n=34) from Eastern EU countries were recorded between 2010 and 2017, encompassing a total of 5567 animals. Hungary was the main exporter to both countries while Poland was the main exporter to France. The highest annual risk of ASF release was related to shipments from Poland to France after 2014 coinciding with an epidemic phase of ASF in that country. During the same period, the risk of ASFV introduction from Hungary to France was much smaller but gradually increased, coinciding with outbreaks in adjacent Ukraine territory during 2015 onward. Despite no outbreaks in France nor Spain were detected to date, quantified values of ASF introduction risk into Western Europe were very high, suggesting that prohibiting international WB trade in 2018 was well justified. Our study highlights the relevance to monitor and supervise more closely legal wildlife production within the EU countries. Our results are an interesting example on how to use trade data to design and implement risk reduction strategies to reduce the spread of pathogens in the wildlife trade sector.

**Highlights:**

⍰ This is the first documented report of the legal trade of live WB between EU countries during the last decade (2010-2017) prior to the EU Commission’s decision to prohibit shipments of live wild pigs from ASF-infected areas in June 2018.
⍰ It provides insight into the level of trade between Eastern and Southern EU countries between 2010 and 2017.
⍰ Results highlight that the risks of disease introduction and dissemination of African swine fever through the legal wildlife trade during the study period were very high and measures taken by banning WB trade in June 2018 were totally justified.
⍰ Our data suggest that collecting and sharing information on wildlife production and trade can be beneficial for identifying and managing disease risks.
⍰ This work served as a good example of a risk assessment exercise with potential applications for other infectious pathogens as well as wildlife production and trade situations such as promoting cross-sector collaboration between animal health, agricultural and wildlife authorities or strengthening the design and implementation of regulations in the surveillance of captive wild animals

## 1. Introduction

Several health crises linked with zoonotic disease emergence in the last decade (COVID-19, SARS Co-V, Ebola) have emphasized the important links existing between global health and wildlife trade and exploitation and the need to manage wildlife trade from a One Health perspective [1]. In the livestock sector, one of the pathogens with the highest devastating impact on animal production, food security and farmer livelihoods is African swine fever (ASF). This virus is currently present in more than 80 countries on the planet, and is possibly considered, the most important animal pandemic of recent times [2], equally affecting pig production but also devastating wild pig populations worldwide [3–4]. ASF spreads in multiple ways such as direct contact susceptible and infected animals or its body fluids, contact with contaminated surfaces (fomites) and/or ingestion of contaminated pork products. Since its introduction from Africa to the Caucasus in 2007, ASF has progressively spread to most countries in East and Central Europe and subsequently to most of Asian countries [5–10]. Therefore, managing the risk of ASF introductions into new territories remains a very relevant challenge, due to the absence of a vaccine, the high persistence of the virus in the environment [7, 9]. Despite it is not able to infect humans, it is considered an important disease from the One Health perspective due to its capacity to remain in natural environments but also to generate economic and nutritional impacts, mental stress and disruption among subsistence farmers [2].

Wild boar (*Sus scrofa*) farming gained popularity across Europe at the end of last century as a sustainable and profitable alternative to traditional livestock as a source of game meat for gourmet markets but also for repopulating hunting estates within the sport hunting industry [11]. A primary motivating factor behind for the development of wild boar (WB) international trade was the considerable size disparity between the WB subspecies from Eastern Europe *Sus scrofa attila* which are notably larger [12], making these animals attractive trophies for Western European hunters. Indeed, an average adult male in the Western EU weighs about 80 kg while Eastern EU specimens can reach up to 250 kg. As a result, legal movements of live WB for the hunting industry became very common within the EU since the end of the last century, but information about this trade has been poorly documented to date. The occurrence of long-distance jumps of the virus through international trade is often linked to human-mediated activities, such as the trade of infected live suids, their meat products or infected materials [10]. Therefore, the design of more efficient ASF prevention and control strategies required a holistic One Health approach that combines i) surveillance systems in wild and domestic pig compartments but also ii) cross-sectorial collaboration between agricultural and veterinary authorities, wildlife managers and the animal trade sector and iii) the implementation of risk management and monitoring tools and regulations. Confronted with the potential challenge of controlling ASF spread incurred by those WB movements [11, 13, 14], the EU Commission decided to prohibit live WB shipments from infected ASF areas in June 2018 [15]. Nevertheless, despite such measures seemed coherent, quantitative estimates on the level of trade occurring prior to this decision were not available. The aim of this study was to implement a risk assessment approach to quantify the levels of WB trade to France and Spain before the ban, identify the likelihood of ASF introduction and spread linked with this activity in the countries concerned and use this information to design safer risk management strategies for trading live WB trade in the EU.

## 2. Materials and Methods

### 2.1 Data sources and movement analysis

Information on declared legal movements of livestock between EU countries is recorded via the Trade Control and Expert System [16], which is a web-based veterinary certification tool. In this study, we analyzed a total of 127 trade movements of live WB entering Spain and France from other EU countries between 2010 and 2017. This quantification was obtained from the TRACES database in order to (i) identify the primary routes of exposure (WB breeding or hunting), and ii) quantify the spatial and temporal variation in risk, including country of origin, country of destination and year. For France, data encompassed 93 movements of WB during the eight-year observation period from 2010 to 2017, whereas for Spain, available data documented 34 movements between 2012 and 2017. The movements reported before 2014 (n=74) only contained details on the species (*Sus scrofa*), year and month of the trade movement, country of origin and destination and the quantity of individuals moved. After 2014, the available information on movements (n=53) was more detailed, including a copy of the movement certificate, the date of declaration of the movement, the method and type of WB identification, the names and addresses of the sender and the recipient, the duration of the transport, and information on veterinary inspections (number of inspections carried out, date, and location).This information allowed us to quantify various factors, including the number of movements, the average quantity of WB per movement, WB reception and departure facilities, the average transport time, the mode and type of identification and the potential inspections carried out.

Welch’s *t*-test was used to compare the average number of movements of various trade routes. The Wilcoxon signed-rank test was performed to compare medians of the total number of WBs transported per year. A statistical test of normality was carried out prior to the Welch test to ensure that the distributions were normal and comparable.

### 2.2. Risk estimation

To quantify the risk of ASF introduction through the trade of live WB, we developed a probabilistic model according to current international WOAH standards in risk assessment [17]. A list of key parameters, assumptions and references for the model are presented in Table 3. The principal output of the model is the probability of exposing susceptible hosts to ASFV in a given destination country following the importation of live WB from a given source country - some authors call this probability the *release* probability. Such probability was estimated annually, using TRACES data of legal trade over the period 2014-2017. The release probability model is expressed as follows:

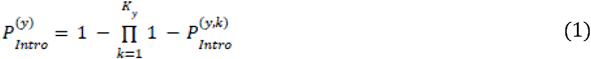

where () is the introduction probability for a given year, (,) is the introduction probability associated with the ^th^ shipment of year, and is the number of shipments from the source country to the destination country during year. The calculation of (,) requires the calculation of two probabilities for each WB concerned (Fig 1, Table 3) that account for chronological events within the *country of origin* (CO) and within the *destination country* (DC). The former quantifies the probability of a transported animal carrying ASFV and releasing the virus from its country of origin; the latter quantifies the probability of the infectious animal participating in a transmission event upon arrival in the destination country. Thus, (,) is calculated as follows:

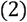

where () and are the country of 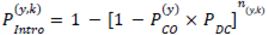 components of the release probability associated with each animal shipped during year, and _(,)_ is the number of WB within the ^th^ shipment of that year.

**Figure 1:**
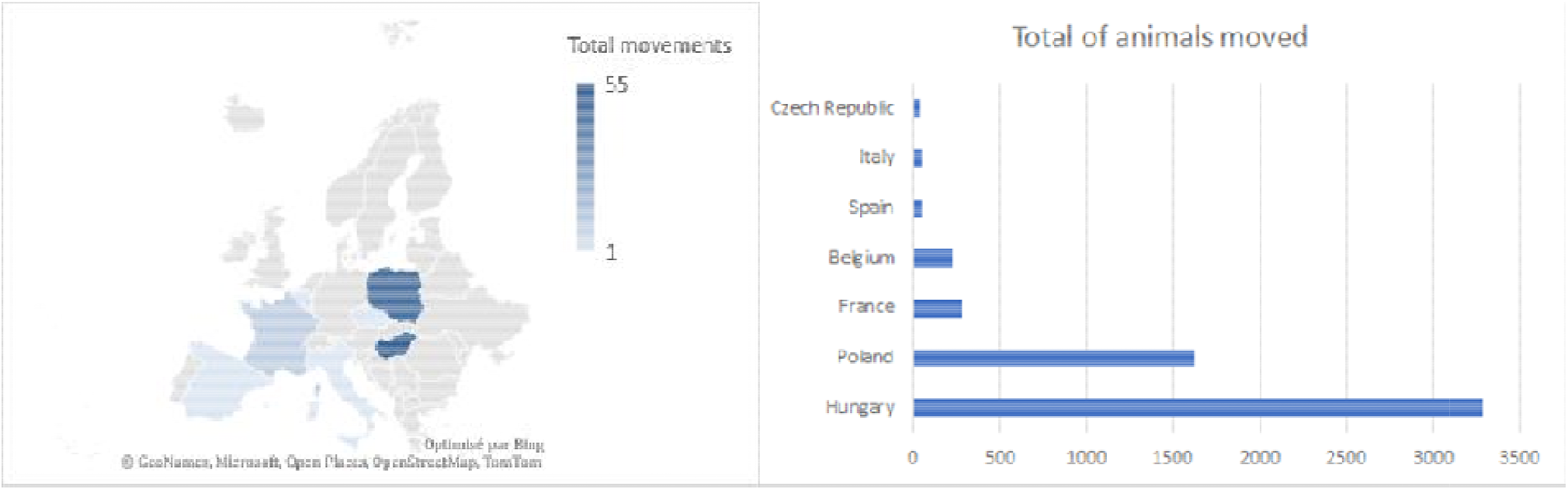
Illustration of shipments and animals of our study. Right: Map of the major countries involved in origin of wild boar trade into France and Spain represented by the number of movements in our study (minimum 1 and maximum 55). Left: Graph of the total of animals moved from each country of origin.

### 2.3. Release probability in the country of origin

Based on previous studies that have quantified the risk of ASF introduction [18–20], we hypothesized that two different mutually compatible and non-exclusive scenarios could potentially lead to release events in the process of a commercial transaction between two hunting farms in different countries. Those are represented in Figure 1:

> Scenario 1: an apparently healthy WB is *incubating* a pre-symptomatic ASFV infection when loaded into a transportation vehicle in the country of origin, meaning the animal becomes *infected prior to transportation*.
>
> Scenario 2: a susceptible WB becomes infected with ASFV following contact with contaminated fomites within the transportation vehicle, meaning the animal becomes *infected during transportation*.

Details on the equations to model these two scenarios can be found in Supplementary Material.

### 2.4. Release probability in the destination country

The conditional probability of release events in the destination country () was defined as the probability that an imported infected WB would come into contact with at least one susceptible individual and transmit the virus to the local WB population. This is the product of two independent events, which are the probability of survival of transportation () and the probability of an outbreak after the introduction of an infected boar () and was calculated as follows:

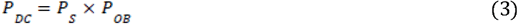

### 2.5. Annual incidence in WB

The annual incidence in WB 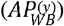 was estimated differently depending on the country of origin. For Poland, the incidence was assessed by a Hidden Markov Model (HMM), based on Polish data on WB population [20], combined with data on WB cases between 2012 and 2017 [20–23]. For Hungary, it was estimated by a kernel estimation (Kernel Density Estimation – KDE), from the probability of detection generated by HMM taking population data from National Game Management Database [23] and data on WB cases from neighboring Ukraine between 2016 and 2017 [24, 26]. Other sources of information for the inputs of the model are available in Supplementary material.

Two important assumptions were made in this approach: i) The annual incidence in both countries was considered the same. ii) Given that Hungary had no official cases until 2018 but there was ASF circulation at the Ukrainian border since late 2016 at less than 80 Km from one of the exporting Hungary farms, we considered these cases and the potential transboundary spillover from Ukraine in our risk calculations. Details on the calculations of the annual incidence of ASF in each country can be found in Supplementary Material

### 2.6. Monte Carlo Simulation

The model was coded and run using R (R Core Team, 2024). Monte Carlo simulation was used to explore the uncertainty in the principal model output 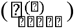 given the uncertainties in each of the model parameters. Analyses were based on the results of 10,000 stochastic simulations.

### 2.7. Sensitivity analysis

Sensitivity analysis was performed to measure the effect of variability and uncertainty in model parameters on the model outcomes. To achieve this, we used the method of Sobol indices [27]. This method associates an output variable Y with a single random input (Xi), provides the sensitivity index Si, defined as follows:

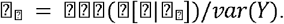

## 3. Results

### 3.1. Quantified number of movements, origin and destination

During the observation period, 127 trade movements (93 to France and 34 to Spain) were reported, encompassing a total of 5567 transported WBs. The average annual number of shipments sent to Western Europe was estimated at 21.3± 6.6 trades/year. From the 57 more detailed movements (2014 to 2017), the average transport time was estimated at 14h50 (median=16h00, max=33h00, min=09h43). Table 1 and Figure 1 summarize trade movements of live WB between the different European countries across the observation period. Hungary was the main country exporting live WB to France and Spain, with a total of 55 trade movements and 3286 animals, while Poland was the main exporter to France in terms of transactions (48 movements and 1617 animals). The rest of the movements occurred mainly between Western EU countries: 12 between France and Spain and 10 between France and Belgium. Only one transaction included Italy and the Czech Republic.

**Table 1:** Table summarizing all incoming wild boar trade movements reported by France and Spain in the TRACES database during the period 2010-2017.

| **Year** | **Movements to**<br>**France** | **Animals to**<br>**France** | **Movements to**<br>**Spain** | **Animals to**<br>**Spain** | **Total of**<br>**movements** | **Total of**<br>**animals** |
| --- | --- | --- | --- | --- | --- | --- |
| **2010** | **1** | **12** | **N/A** | **N/A** | **1** | **12** |
| **2011** | **1** | **21** | **N/A** | **N/A** | **1** | **21** |
| **2012** | **10** | **199** | **3** | **90** | **13** | **289** |
| **2013** | **20** | **800** | **1** | **26** | **21** | **826** |
| **2014** | **27** | **1423** | **3** | **46** | **30** | **1469** |
| **2015** | **16** | **755** | **5** | **281** | **21** | **1036** |
| **2016** | **14** | **602** | **12** | **546** | **26** | **1148** |
| **2017** | **4** | **295** | **10** | **471** | **14** | **766** |
| **TOTAL** | **93** | **4107** | **34** | **1460** | **127** | **5567** |

Between 2010 and 2017, the median number of animals transported per movement was estimated at 33 WB per movement (max=200, min=1). During the hunting season (October to February), the average quantity of WB exported was significantly higher than that during the rest of the year (p<0.05). Among 57 movements studied, all the WBs transported were identified either individually (75% of cases), or rest by batch. The different modes of individual identification used were ear tags (68%), electronic chips (23%) and bracelets (9%). Only 5 movements (8.7%) reported the implementation of a veterinary check at a median of 14 days after the arrival of the animals at destination, including two movements to France and two to Spain and one to Slovakia.

### 3.2. Specific details for each country

All shipments from Poland originated from a unique wildlife farm located in the West of the country located approximately 400 km from the closest ASF outbreak declared in Poland between 2014 and 2018. This farm was not registered as a WB breeding farm and the origin of the animals could not be identified by local sources. In the case of Hungary, animals originated from 8 different farms located throughout the country. Although ASF was declared in Hungary in 2018, one of the exporting farms was located 100 km from ongoing ASF outbreaks located in Ukraine between 2013 and 2017. In France, all shipments arrived mainly at four farms located in the Departments of Côte-d’Or, Marne and Oise. In Spain, shipments of destination were all located in the center and east of the country (Provinces of Madrid, Valencia and Castilla-La Mancha).

### 3.3. Overall probability of introduction

The annual probability of ASF introduction was calculated selectively for movements between countries in the eastern EU (Hungary and Poland) and their destination in Western EU (France and Spain) between 2012 and 2017. Annual values of risk are shown in Table 2 and Figure 3. The median annual risk for both countries in Southern Europe increased exponentially during the study period, passing from 1.97E-5 in 2012 to 8.10E-3 in 2017 (a 410-fold increase) for Poland-France movements and from 6.14E-4 in 2015 to 1.23E-3 (a two-fold increase) in 2017 for Hungary-Spain movements. The median annual risk over the study period arising from Polish shipments (8.10E-3) was 8.6-fold higher than the same value related to shipments from Hungary (9.37E-4). As a result, since Poland was only exporting to France, the overall annual risk of ASF introduction was clearly higher for France compared to Spain. When comparing annual risks of ASF introduction for Hungarian shipments during 2015-2017, values were 2.9-fold higher for Spain than for France (3.26E-4 for France vs 9.37E-4 for shipments to Spain). The risk due to shipments between Hungary and France peaked in 2014, and then dropped due to a decrease in the number of recorded shipments.

**Table 2:**
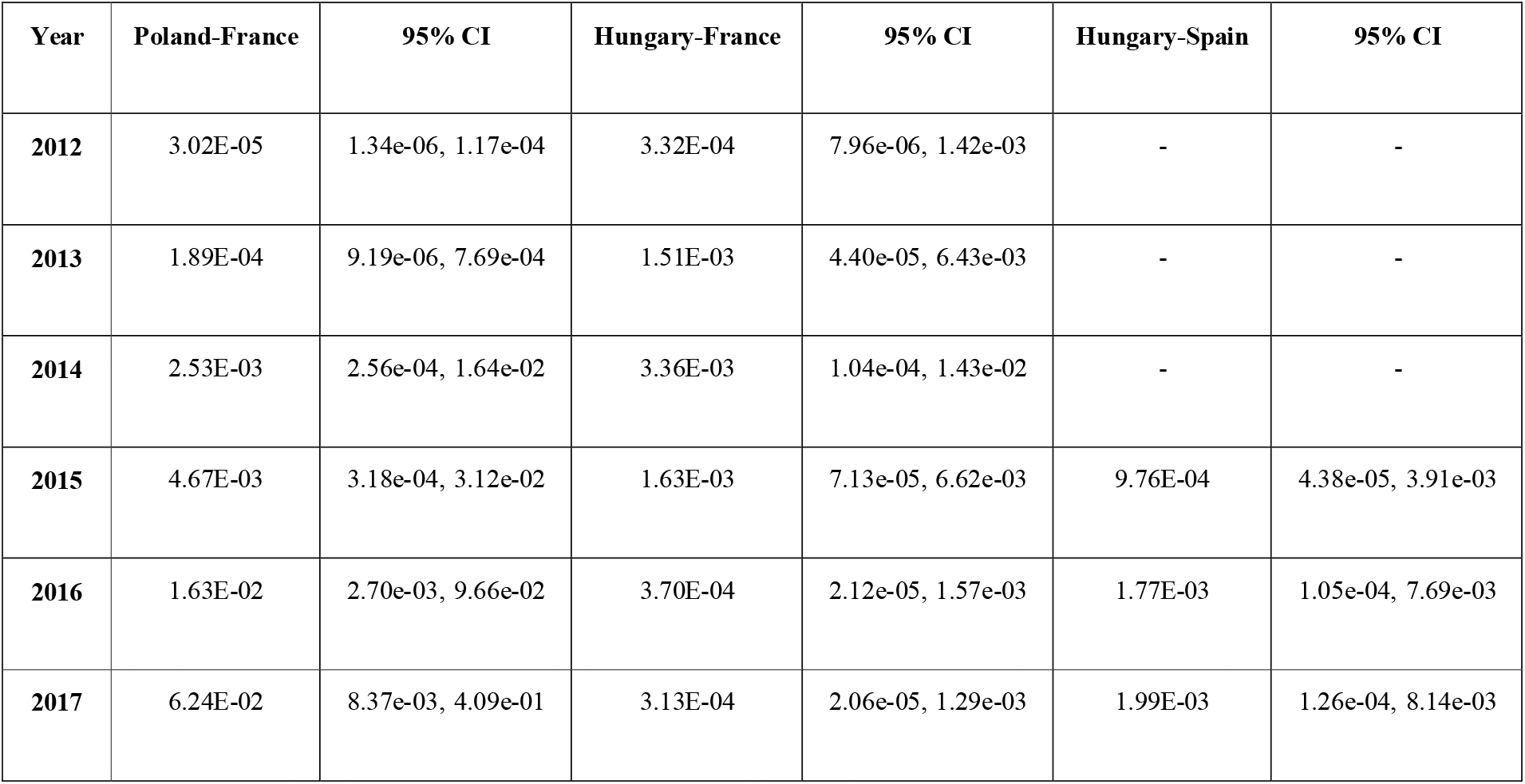
Annual probability of ASF introduction for movements between EU countries between 2015 and 2017.

| **Year** | **Poland-France** | **95% CI** | **Hungary-France** | **95% CI** | **Hungary-Spain** | **95% CI** |
| --- | --- | --- | --- | --- | --- | --- |
| **2012** | 3.02E-05 | 1.34e-06, 1.17e-04 | 3.32E-04 | 7.96e-06, 1.42e-03 | - | - |
| **2013** | 1.89E-04 | 9.19e-06, 7.69e-04 | 1.51E-03 | 4.40e-05, 6.43e-03 | - | - |
| **2014** | 2.53E-03 | 2.56e-04, 1.64e-02 | 3.36E-03 | 1.04e-04, 1.43e-02 | - | - |
| **2015** | 4.67E-03 | 3.18e-04, 3.12e-02 | 1.63E-03 | 7.13e-05, 6.62e-03 | 9.76E-04 | 4.38e-05, 3.91e-03 |
| **2016** | 1.63E-02 | 2.70e-03, 9.66e-02 | 3.70E-04 | 2.12e-05, 1.57e-03 | 1.77E-03 | 1.05e-04, 7.69e-03 |
| **2017** | 6.24E-02 | 8.37e-03, 4.09e-01 | 3.13E-04 | 2.06e-05, 1.29e-03 | 1.99E-03 | 1.26e-04, 8.14e-03 |

**Figure 2:**
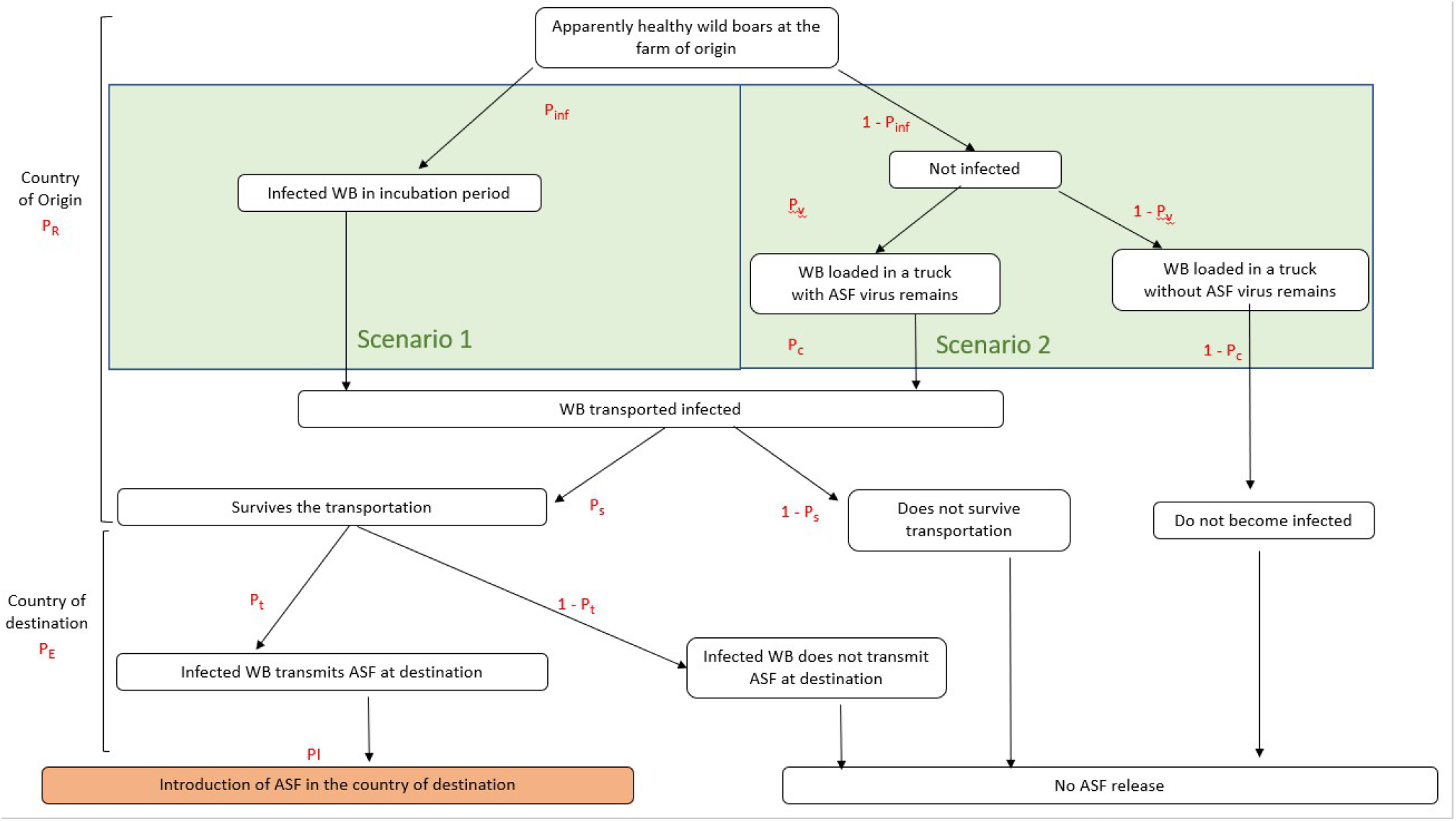
Scenario tree illustrating the different events in the pathway of ASF release which includes events in the country of origin and the country of destination. There two scenarios leading to a situation of wild boars being transported infected: Scenario 1 which considers that wild boars (WB) are loaded in the transport truck during the incubation period and Scenario 2, which considers that wild boars are loaded healthy and become infected with ASF remains in the transport truck

**Figure 3:**
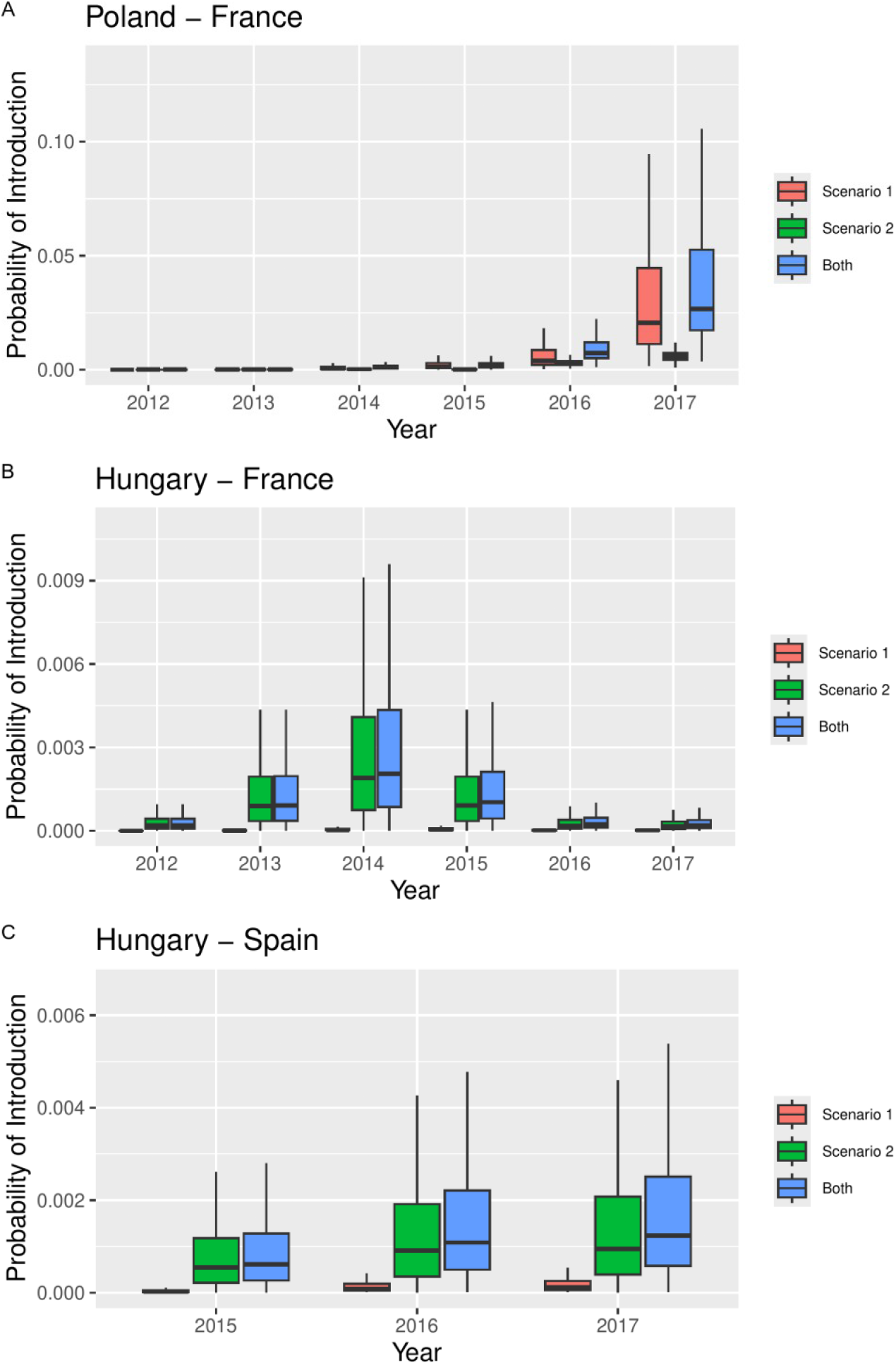
Box plots of the probability of introduction (PI) as a function of year and scenario for movements between A) Poland and France, B) Hungary and France and C) Hungary and Spain, represented on the natural scale on the left and on a logarithmic scale on the right. Scenario1 is describes a situation where apparently healthy wild boars are loaded in the transport truck during the incubation period of ASF and prior to the occurrence of symptoms. This is more likely to occur in the context of Poland shipments, because an important proportion of the animals was suspected to be captured from the wild before being loaded for transport. Scenario 2 represents a situation where wild boar are loaded healthy in the transport truck that is inappropriately disinfected and some animals are able to become infected with contaminated fomites during transport. This scenario appears more likely in the case of Hungarian shipments.

Comparing the risks for scenarios 1 and 2, these were fairly similar in the case of movements from Poland, the risk for scenario 1 being 4.1-fold higher than for scenario 2 during the 2012-2017 period. However, in the case of movements from Hungary, the risk from scenario 2 was 26-fold and 11-fold higher than for scenario 1 for movements to France and Spain respectively.

### 3.4. Sensitivity analysis

Sensitivity analyses identified the parameters with the greatest influence on the model outcomes for each exporting-importing country combination. For each combination of source and destination country of interest, the five highest Sobol indices and their associated parameters or interactions are reported in Table 3.

**Table 3:**
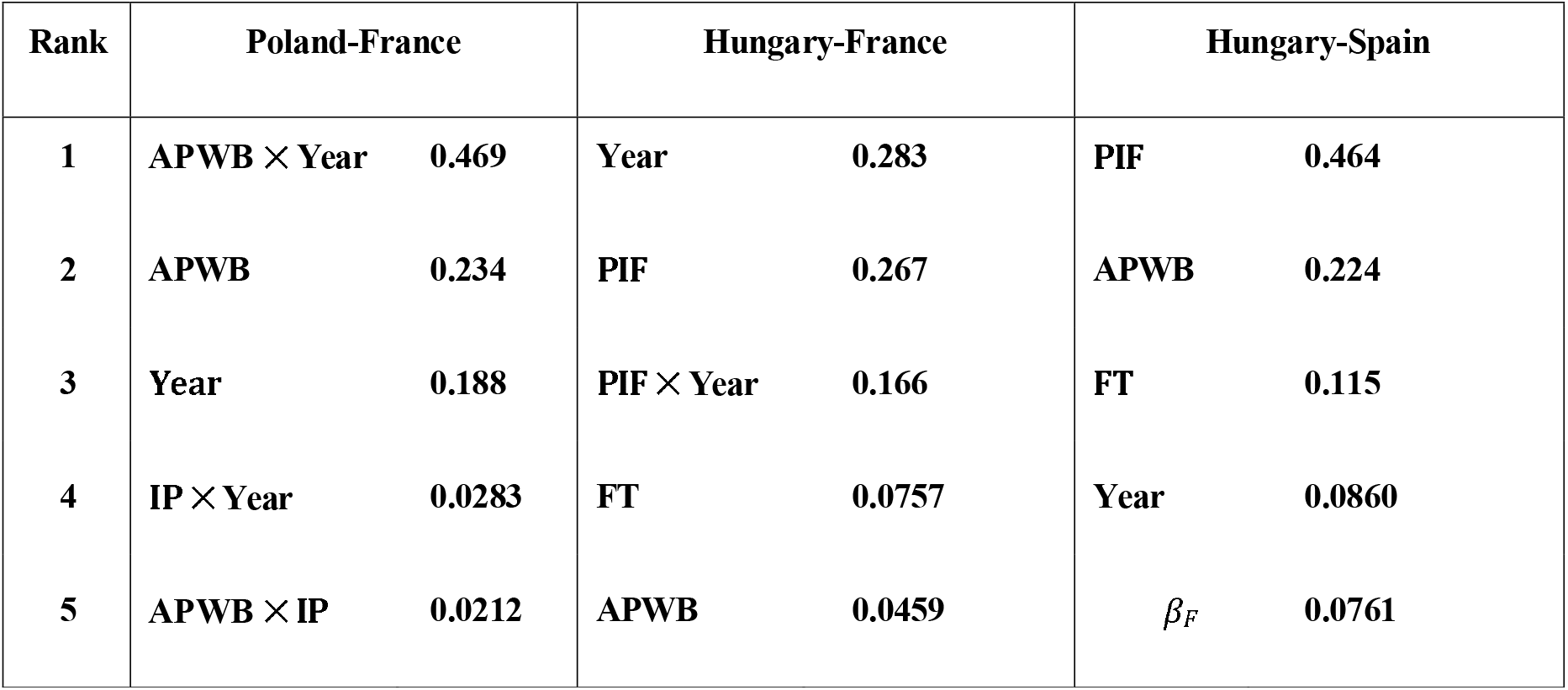
Five largest Sobol indices associated with the sensitivity of PIntro(y) to parameters and two-way parameter interactions, as a function of country of origin and destination. Recall: APWB is the annual proportion of Wild Boar contracting ASF infections; IP is the incubation period; PIF is the proportion of farms infected with ASF in a year; FT is the total number of farms in the country of origin; *β*_*F*_ is the fomites-to-animal transmission rate.

## 4. Discussion

Several studies have quantified the risk of introducing ASF virus into new territories through the trade of domestic pigs or their products [18–20, 28] or natural movements of WB [6,29]. However, this is the first study addressing the transboundary ASF spread through commercial WB movements. Although information available was sometimes incomplete (identification methods, veterinary controls) and the accuracy varied across time periods and countries, the TRACES database provided a unique and reliable source to analyze WB trade over almost a decade. One of the main sources of uncertainty in our model concerned the context of WB farming practices in the countries of origin. Despite our efforts to obtain additional clarity in some of those practices, we were not successful in gathering additional information. However, the available data was sufficient to provide a broad overview of the trade process and a preliminary model that can be further refined if new data became available. Our results indicate that, even when conducted within a legal framework, the EU WB trade in the last decade has posed a significant risk of introducing pathogens into new territories, potentially facilitating long-distance jumps of the ASF virus between countries. A clear seasonal pattern was observed, aligned with the hunting season, suggesting that this period should be prioritized for the implementation of preventive and control measures.

From the One Health perspective, these results have contributed to better understand the complex pathways and connections that can occur between free-ranging populations of WB, captive WB and the potential impact of different WB management practices in different countries on the spread of a pathogen with important consequences in environmental health a wide and devastating socio-economic impact.

Regarding the results of our risk assessment, the highest annual probability of ASF introduction arose from WB trade from Poland to France between 2015 and 2017 which posed an 8.6-fold greater risk than shipments from Hungary (Table 2, Figure 3). This difference was consistent with the higher level of ASF spread reported in Poland compared to Hungary during the study period. Despite a decrease in Polish WB shipments after their official declaration of first ASF cases in 2014 [22,23, 30] an average of 15 shipments and 750 animals per year continued arriving annually to France. Despite the first official ASF cases in Hungary were only declared in 2018 [5,7], we considered that risk values could not be negligible on the basis of the high number of shipments involved (n=53) combined with the proximity of a WB farm located at only 70 km from active Ukrainian ASF outbreaks reported between 2015 and 2017.

In addition, we found different WB management situations and stock management strategies in the countries of origin which were likely to influence risk outputs from the model. Because the origins of WB from Poland were unclear and could have occasionally been sourced from free ranging populations, we assumed this situation could have influenced the probability of loading WB with subclinical ASF infection, which explains the higher level of risk for Scenario 1 (Figure 3). Conversely, because Hungarian shipments came from well-established WB farms which were raised similarly to domestic pigs, we assumed a better capacity to detect and prevent loading animals with a latent ASF infection (Figure 3).

There is an assumption that legal wildlife trade is done under veterinary supervision. However, our data suggest that veterinary control inspections were scarce (8.7% of shipments) in all the countries concerned, highlighting the challenges to implement disease surveillance in wildlife species [11, 13, 14] and the need to increase wildlife management capacities among veterinary authorities [30, 31].

Our model primarily focused on the risk of ASF release at the country of origin, whereas potential exposure events in the destination countries were oversimplified. We assumed that the life span of WB released in the country of destination was short and that contacts with other wild or domestic pigs were highly unlikely. Further studies are needed to consider additional exposure scenarios which could vary depending on the geographic characteristics of the release location. While France and Spain still remain free from the disease in 2025, the increasingly high-risk values of our model since 2014 suggest the possibility that ASF virus could have been introduced without generating detectable outbreaks.

Our risk output models indicate that the ban on international WB movements between EU countries [15], implemented in June 2018 was timely and justified. However, forbidden transactions can often generate the development of alternative illegal markets [32]. Coincidentally, an illegal WB introduction was pointed out as one of the drivers of the unexpected introduction of ASF into Belgium 3 months after the ban [33, 34], although evidence for this hypothesis could never be proven.

Currently, the continuous spread of ASF across the EU is leading to significant changes in WB management activities, including the likely reduction or elimination of operational WB farms in ASF affected countries. Despite trade information collected in our study was retrospective and the current epidemiological and WB production context has changed, our risk assessment approach represents a good conceptual model, adaptable for further applications to other wildlife species, pathogens or trade situations.

## Conclusion

Our study illustrated the challenges of monitoring wildlife production, even in developed countries operating under a legal framework and the advantages of collecting and quantifying data on wildlife trade shipments for the purpose of monitoring the risk of a highly relevant transboundary pathogen dissemination such as ASF. Our results highlight the need to manage wildlife trade from a holistic One Health perspective. such as enhancing cross sector collaboration between wildlife management and veterinary authorities, promoting higher levels of disease awareness trade stakeholders or the implementation and enforcement of harmonized regulations within the legal wildlife trade in the EU.

## Ethics statement

Not applicable

## Funding

This research was partially co-funded by the COST Action CA15116, ASF-STOP supported by COST (European Cooperation in Science and Technology), by the European Union’s Horizon Europe Project 10113646 EUPAHW and by the Partnership between CIRAD and DGAL in France and Jose Angel Barasona is a recipient of a “Ramón y Cajal” contract [RYC2022-038060-I] funded by the Spanish Ministry of Science and Innovation (MCIN/AEI) and Fondo Social Europeo Plus (FSE+).Views and opinions expressed are those of the author(s) only and do not necessarily reflect those of the European Union or the European Research Executive Agency.

Neither the European Union nor the granting authority can be held responsible for them

## CRediT authorship contribution statement

**Jori, F**., Conceptualization, Methodology, Investigation, Data curation, Formal analysis, Investigation, Visualization, Validation, Writing – original draft, Writing – review & editing, Project administration, Supervision, Funding acquisition

**Pleydell, D.R.J**., Data curation, Formal analysis, Data modeling and Visualization, Validation, Writing – original draft, Writing – review & editing

**Burnichon, E**., Methodology, Formal analysis, Visualization, Data curation, Investigation, Validation, Writing.

**Casal, J**., Methodology, Investigation, Writing – review & editing.

**Barasona, J.A**., Conceptualization, Methodology, Visualization, Validation, Writing – review & editing,

## Data availability

The authors do not have the permission to share the raw data.

## Acknowledgements

We would like to acknowledge several contributors who provided instrumental information to facilitate this work in different EU countries including Andrzej Jarynowsky, András Náhlik, Henryk Okarma, Jacques Vion and Beatriz Martinez Lopez. We thank Germán Caceres and Agnès Giraud for facilitating access to TRACES information.

