## Supplementary material for "Wild boar trade and African swine fever risk of introduction into new territories: A quantitative release assessment with retrospective data of wild boar shipments to France and Spain (2010-2017)"

1. ***Risk estimation model***

Here we present in full the probabilistic model that was developed to quantify the risk of ASF introduction through the trade of live wild boar. A list of key parameters, assumptions and references for the model are presented in Table 3. The principal output of the model is the probability of exposing susceptible hosts to ASFV in a given destination country following the importation of live wild boar from a given source country - some authors call this probability the *release* probability. This probability was estimated annually, using TRACES data of legal trade over the period 2014-2017. The probability model is expressed as follows:

$P_{Intro}^{(y)}=1-\prod_{k=1}^{K_{y}} 1-P_{Intro}^{(y,k)}$	(1)

where $P_{Intro}^{(y)}$ is the introduction probability for a given year $y$, $P_{Intro}^{(y,k)}$ is the introduction probability associated with the $k$^th^ shipment of year $y$, and $K_{y}$ is the number of shipments from the source country to the destination country during year $y$. The calculation of $P_{Intro}^{(y,k)}$ requires the calculation of two probabilities for each wild boar concerned (Fig 1, Table 3, main paper) that account for chronological events within the *country of origin* (CO) and within the *destination country* (DC). The former quantifies the probability of a transported animal carrying ASFV and releasing the virus from its country of origin; the latter quantifies the probability of the infectious animal participating in a transmission event upon arrival in the destination country. Thus, $P_{Intro}^{(y,k)}$ is calculated as follows:

$P_{Intro}^{(y,k)}=1-{[1-P_{CO}^{(y)}\times P_{DC}]}^{n_{(y,k)}}$	(2)

where $P_{CO}^{(y)}$ and $P_{DC}$ are the country of origin and destination country components of the release probability associated with each animal shipped during year $y$, and $n_{(y,k)}$ is the number of wild boar within the $k$^th^ shipment of that year.

1.1. ***Release probability in the country of origin***

We hypothesized that two different mutually compatible and non-exclusive scenarios could potentially lead to release from the country of origin event following a commercial transaction between two hunting farms in different countries:

Scenario 1: an apparently healthy wild boar is *incubating* a pre-symptomatic ASFV infection when loaded into a transportation vehicle in the country of origin, meaning the animal becomes *infected prior to transportation*.

Scenario 2: a susceptible wild boar becomes infected with ASFV following contact with contaminated fomites within the transportation vehicle, meaning the animal becomes *infected during transportation*. Details on the equations to model these two scenarios can be found in Supplementary Material.

We assume throughout that wild boar displaying symptoms would not be transported, due to their reduced market value. Let $P_{Inc}^{(y)}$ and $P_{Trans}^{(y)}$ represent the probabilities of these two scenarios respectively, for a given year $y$. The CO release probability is calculated as the complement to the probability that neither scenario occurs, which, assuming independence, is the given by

$P_{CO}^{(y)}=1-(1-P_{Inc}^{(y)})(1-P_{Trans}^{(y)})$	(3)

This probabilistic approach is similar to that of previous authors who have quantified the risk of African swine fever introduction (Mur et al., 2012).

1.1.1. ***Scenario 1: wild boar infected prior to transportation***

We assume that every wild boar involved in international trade would appear healthy at the time of capture and transportation, meaning that an infection would need to be latent for an infected wild boar to be loaded onto a transportation vehicle. Moreover, we assume that this disease status has no effect on the probability of being captured. The probability that a wild boar is infected prior to transportation is therefore assumed to be

$P_{Inc}^{(y)}={AP}_{WB}^{(y)}\times\frac{IP}{365}$	(4)

where ${AP}_{WB}^{(y)}$ is the annual proportion of wild boar becoming infected with ASFV in a source country during year *y*, $IP$ is the duration of the incubation period in days and $\frac{IP}{365}$ approximates the proportion of the year covered by a typical incubation period. The estimation of ${AP}_{WB}^{(y)}$ for the two potential source countries in our study is detailed below.

1.1.2. ***Scenario 2: wild boar infected during transportation***

The probability that a healthy wild boar is infected by fomites within a contaminated transportation vehicle is modeled as

$P_{Trans}^{(y)}=P_{V}^{(y)}P_{C}$	(5)

where $P_{V}^{(y)}$ is the probability that a vehicle is contaminated, and $P_{C}$ is the probability that ASFV transmits from a contaminated vehicle to a susceptible wild boar during transit. These two probabilities are modeled as follows

$P_{C}=1-exp\left( -\frac{\beta t_{V}}{2} \right)$	(6)

and

$P_{V}^{(y)}={AP}_{IF}^{(y)}\times P_{VID}$	(7)

where ${AP}_{IF}^{(y)}$ is the annual proportion of pig farms declaring ASF infections in year *y* in the source country, which is used as a proxy for the prevalence of contamination among livestock transportation vehicles, and $P_{VID}$ is the probability that a vehicle is insufficiently disinfected. The fomites to wild boar transmission rate was assumed to be half of published estimates of pig to pig transmission rates: in other words, the probability for a wild boar becoming infected when in contact with infected fomites in a truck was considered half that of a healthy pig becoming infected when coming into contact with an ASFV infected pig. In the above formula $\beta$ represents the daily rate of transmission between two pigs, determined experimentally (Vergne et al. 2021) and represented by a Pert(0.2, 0.4, 0.6) distribution. The parameter $t_{V}$ represents the expected transport time in days, which was estimated at 24 hr divided by the median transport time of 16 hours reported in the TRACES database. We assumed that the probability of a healthy and susceptible infected pig becoming infected with ASF after spending a median time of 16 hours in an inadequately disinfected transport vehicle was the same as becoming infected when in contact with an excreting infected pig (Vergne *et a*l. 2012). The probability for a pig transport vehicle being inefficiently cleaned and disinfected ($P_{VID}$) was obtained from a study implemented in Germany (Weber and Memmken, 2018) and the assumption that the context would be similar in Hungary and Poland. Survival probabilities for wild boars during transport were estimated based on expert opinion data from veterinarians with experience in wild boar farming in France and Spain. Their opinions were converted into Pert distributions and combined into a mixture distribution (Table 1).

1.2. ***Release probability in the destination country***

The conditional probability of release events in the destination country ($P_{DC}$) was defined as the probability that an imported infected wild boar would come into contact with at least one susceptible individual and transmit the virus to the local wild boar population. This is the product of two independent events, which are the probability of survival of transportation ($P_{S}$) and the probability of an outbreak after the introduction of an infected boar ($P_{OB}$) and was calculated as follows:

$P_{DC}=P_{S}\times P_{OB}$	(8)

The average time of transport was estimated on the basis of the initial and final transaction dates available in the TRACES database and in all cases was less than 24h. This journey duration was considered too short for wild boars loaded during the incubation period, to die during transport because of ASF infection. The probability for a wild boar to survive to road transport by truck ($P_{S}$) was estimated by the combination of two Pert distributions, obtained from expert opinion of two veterinarians working in hunting farms and dealing with reception of wild boar shipments. The probability that wild boar would die after such a short journey was considered negligible and the reliability of their response was estimated between 80% and 90%.

The probability of an outbreak occurring in the arrival facility after the introduction of a wild boar Infected ($P_{t}$) was obtained from an experimental infection work of Guinat et al. (2016). This work considers that the probability of an outbreak not occurring in a pig population following the introduction of an ASF virus infected pig ($P_{NOB}$) can be represented by a Pert distribution ranging between 0.07 and 0.16 (most likely 0.10). The inverse probability $P_{t}$ (probability that an animal susceptible becomes infected following the introduction of an infected individual) has therefore been calculated. In this case, any boar arriving at destination of hunting or breeding establishment is supposed to survive long enough (minimum 6 days) to shed the virus and transmit it – this number of days is defined according to the average length of the period latency (period before excretion begins) experimentally determined between 5.8 and 9.7 (Guinat et al.,2018).

1.3. ***Annual incidence in wild boar***

The annual incidence in wild boar (${AP}_{WB}^{(y)}$) was estimated differently depending on the country of origin. For Poland, the incidence was assessed by a Hidden Markov Model (HMM), based on Polish data on wild boar population (Statistics Poland, 2021) combined with data on wild boars cases between 2012 and 2017 (ADNS, 2021). For Hungary, it was estimated by a kernel estimation (Kernel Density Estimation – KDE), from the probability of detection generated by HMM taking population data from National Game Management Database (1996) and data on wild boar cases from neighboring Ukraine between 2016 and 2017 (ADNS, 2021 and EMPRES-i, 2021).

Two important assumptions were made in this approach: i) The annual incidence in both countries was considered the same. ii) Given that Hungary had no official cases until 2018 but there was ASF circulation at the Ukrainian border since late 2016 at less than 80 Km from one of the exporting Hungary farms, we considered these cases and the potential transboundary spillover from Ukraine in our risk calculations.

1.3.1. ***Annual incidence proportion in wild boar: Poland***

A hidden Markov model was used to model the growth of the annual incidence ($I$) of ASF in Polish wild boar over the study period. In this model the unobserved stochastic dynamics of annual incidence was modeled as the following auto-correlative process

$${I(t)}\sim Lnorm(ln((1+Ɣ) \times I(t-1) + \epsilon_{Imp}),\sigma_{I})$$

Where ${I(t)}$is the total number of ASF cases in year *t*, 1+$Ɣ$ is the annual growth rate, $\epsilon_{Imp}$ is the annual importation rate, and $\sigma_{I}$ is a standard deviation term which accounts for stochastic variation in $I(t)$. For simplicity, all parameters in this model are assumed constant in space and time. The incidence proportion (${AP}_{WB}(t)$) was obtained as

$${AP}_{WB}(t)=\frac{I(t)}{WB(t)}$$

where $WB(t)$ is the official Polish estimate of the wild boar density in year $t$, where available, or an estimate imputed with the model

$${WB(t) \sim Norm(\mu_{WB},\sigma_{WB}).}$$

The observed incidence was assumed to arise as follows:

$$I_{Ob}(t) \sim Poisson(p_{Det}I(t)+\epsilon)$$

where $p_{Det}$ is the probability to detect a given ASF case in wild boar, and the tolerance parameter $\epsilon$ was set to ${10}^{-11}$.

This model was parameterised for years 2010-2017, and we set the following initial condition:

$I(t=2010)=0$,

${AP}_{WB}(t=2010)=0$.

The model was specified in the Baysian paradigm, with the following priors:

$Ɣ\sim Exp(1)$

$$\epsilon_{Imp} \sim Exp(1)$$

$$\sigma_{S} \sim Exp(rate={10}^{-1})$$

$\mu_{WB} \sim Exp(rate={10}^{-6})$

$$\sigma_{WB} \sim Exp(rate={10}^{-6})$$

$p_{Det} \sim$Beta(1,1)

The model was specified in Nimble (de Valpine *et al*, 2016), with priors specified using the nimbleNoBounds package (Pleydell, 2024) to aid computational efficiency. The posterior distribution of the parameters was approximated via Markov chain Monte Carlo (MCMC): more specifically, using a combination of Nimble’s block and univariate Metropolis-Hastings samplers. Our MCMC was run for a burn-in period of ${1.1\times10}^{5}$ iterations, followed by a further ${1.1\times10}^{9}$ iterations with thinning set to ${1.1\times10}^{5}$ iterations, resulting in ${10}^{4}$ MCMC samples.

1.3.2. ***Annual incidence proportion in wild boar: Hungary***

There were no official reports of ASF infected wild boar from Hungary during the study period. However, there were official reports of cases very close to Hungary in the Ukraine. Given this lack of data we estimated the incidence proportion (${AP}_{WB}$) of infected wild boar in Hungary as follows. For the years 2016 and 2017, the density of infected wild boar was estimated across Hungary using kernel density estimation to extrapolate the Ukrainian case data. For each of these two years this estimate of the density of infected wild boar was divided by MCMC estimates for the detection probability $p_{Det}$- to correct for under detection - and normalized by the official total wild boar density census estimate, to obtain an estimate of the required incidence proportion (${AP}_{WB}^{(y)}$). For earlier years, the annual incidence was estimated assuming that the rate of growth of the epidemic in Hungary matched the expected value of the growth rate estimated for Poland using the hidden markov model, i.e.

${AP}_{WB}^{(y-1)} = {AP}_{WB}^{(y)} / E[Ɣ]$.

1.4. ***Sensitivity analysis***

Sensitivity analysis was performed to measure the effect of variability and uncertainty in model parameters on the model outcomes. For this we used the method of Sobol indices (Sobol, 2001; Chastaing et al., 2013). This method uses a formula associating an output variable *Y* to a single random input (*X_i_*), and provides the sensitivity index *S_i_* is defined as follows:

$$S_{i}=var(E[Y|X_{i}])/var(Y)$$

These sensitivity indices are based on a decomposition of the overall variance of the model. They allow the calculation of the total contribution of each input factor/parameter to the variance of the output variable. The total contribution of a factor includes the main effect of that factor as well as the interactions involving that factor, i.e. interactions with other factors. In our case the output parameters $(Y)$ of interest were the $P_{Intro}$ estimated for each combination of source and destination country for which the TRACES data was available. The parameters tested in the sensitivity analysis were: annual incidence proportion in wild boar (${AP}_{WB}^{(y)}$); total number of pig farms ($F_{T}$); ASF incubation period ($IP$); annual incidence of infected pig farms (${AP}_{IF}^{(y)}$); the probability that vehicles are insufficiently disinfected ($P_{VID}$); transmission rate ($\beta$); probability that an infected wild boar survives transportation ($P_{S}$); the probability of no outbreak at destination farm following the importation of an ASF infected wild boar ($P_{NOB}$); and the year ($y$).

**Table 3:** Summary of input variables used to calculate the risk of introducing ASFV into France and Spain through the trade of live wild boar.

| **Notation** | **Definition** | **Source** | **Parametrization** |
| --- | --- | --- | --- |
| **Release from country of origin (CO)**  (ASFV leaves country of origin) | | | |
| PCO(y) | Probability of release (i.e. at least one infected wild boar is imported into the destination country) in year *y* |  | PCO(y)=1-(1-PInc(y))(1-PTrans(y)) |
| **Scenario 1: wild boar infected prior to transportation**  (Captured and transported during incubation period) | | | |
| PInc(y) | Probability that an apparently healthy wild boar is incubating an infection when captured |  | PInc(y)=APWB(y) × IP / 365 |
| APWB(y) | Annual proportion of wild boar infected with ASFV in  year *y* |  | Hidden Markov model (Poland),  Kernel density estimation (Hungary) |
| IP | Duration of incubation period (in days) | [6] | Uniform (1, 10) |
| **Scenario 2: wild boar infected during transportation**  (Infected by a contaminated transportation vehicle) | | | |
| PTrans(y) | Probability a healthy wild boar is infected by an insufficiently disinfected transport vehicle |  | PTrans(y)=PV(y)×PC |
| PV(y) | Probability to load a wild boar in an contaminated transport vehicle |  | PV(y)=APIF(y)PVID |
| APIF(y) | Annual proportion of farms with infected pigs, year *y* |  | Beta (FI(y)+1,FT-FI(y)+1) |
| FI(y) | Number of outbreaks on pig farms in year *y* | [7] |  |
| FT | Total number of pig farms |  | Log Normal(μ,σ) |
| PFI(y) | Proportion of infected farms |  | FI(y)/FT |
| , | Mean & st.dev. from pig farms reported in years 2012-2017 | [8] |  |
| PVID | Probability the transport vehicle is insufficiently disinfected or cleaned | [4] | Beta(17.9,18.5) |
| PC | Probability of viral transmission from a contaminated vehicle to transported wild boar |  | PC=1-exp-β × tV2 |
|  | Daily transmission rate between pigs | [9] | Pert (0.2,0.4,0.6) |
| tV | Duration of transportation (in days) | [10] | tV=1 |
| **RELEASE AT THE DESTINATION COUNTRY (DC)**  (Transmission of ASFV in destination country) | | | |
| PDC | Probability of release within destination country. At least one infected wild boar contacts with a susceptible animal & transmits ASFV in destination country |  | PDC=PS×POB |
| PS | Probability of infected wild boar surviving transport | Expert opinion of wild boar managers | Mixture (Pert (0.70, 0.85, 1), Pert (0.80, 0.95, 1)) |
| POB | Prob. of outbreak after introduction of infected wild boar |  | POB=1-PNOB |
| PNOB | Prob. no outbreak after infected wild boar introduced at farm | [5] | Pert (0.07, 0.10, 0.16) |
| **INTRODUCTION RISK**  (Integration with official trade data) | | | |
| PIntro(y,k) | Proba. of introduction via *k*^th^ wild boar shipment of year *y* |  | PIntro(y,k)=1 - 1 -PCO(y)PDCN(y,k) |
| N(y,k) | Number of wild boar transported in the *k*^th^ shipment of year *y* | [10] |  |
| K(y) | Number of wild boar shipments in year *y* | [10] |  |
| PIntro(y) | Probability of introduction in year *y*: i.e. introduction of ASFV into destination country via the international transfer of live wild boar |  | PIntro(y)=1-k=1K(y)1-PIntro(y,k) |
